# *DYNC1H1* mutations associated with neurological diseases compromise processivity of dynein-dynactin-cargo adaptor complexes

**DOI:** 10.1101/092791

**Authors:** Ha Thi Hoang, Max A. Schlager, Andrew P. Carter, Simon L Bullock

## Abstract

Mutations in the human *DYNC1H1* gene are associated with neurological diseases. *DYNC1H1* encodes the heavy chain of cytoplasmic dynein-1, a 1.4 MDa motor complex that traffics organelles, vesicles and macromolecules towards microtubule minus ends. The effects of the *DYNC1H1* mutations on dynein motility, and consequently their links to neuropathology, are not understood. Here, we address this issue using a recombinant expression system for human dynein coupled to single-molecule resolution *in vitro* motility assays. We functionally characterise 14 *DYNC1H1* mutations identified in humans diagnosed with malformations in cortical development (MCD) or spinal muscular atrophy with lower extremity predominance (SMALED), as well as three mutations that cause motor and sensory defects in mice. Two of the human mutations, R1962C and H3822P, strongly interfere with dynein’s core mechanochemical properties. The remaining mutations selectively compromise the processive mode of dynein movement that is activated by binding to the accessory complex dynactin and the cargo adaptor BICD2. Mutations with the strongest effects on dynein motility *in vitro* are associated with MCD. The vast majority of mutations do not affect binding of dynein to dynactin and BICD2, and are therefore expected to result in linkage of cargoes to dynein-dynactin complexes that have defective long-range motility. This observation offers an explanation for the dominant effects of *DYNC1H1* mutations *in vivo*. Collectively, our results suggest that compromised processivity of cargo-motor assemblies contributes to human neurological disease and provide insight into the influence of different regions of the heavy chain on dynein motility.

## INTRODUCTION

Microtubule motors play a key role in the sorting of many cellular constituents, including organelles, vesicles, aggregated proteins and macromolecules. Because of their elongated processes, neurons are particularly reliant on a fully operational microtubule-based transport system. This point is underscored by the association of several mutations in microtubule motors and their co-factors with human neurological diseases (1–6). How these mutations affect transport mechanisms is poorly understood. Addressing this issue is expected to provide insight into the cellular basis of neurological disease, and also inform diagnostic and therapeutic efforts.

Mutations in the gene encoding the heavy chain of the cytoplasmic dynein-1 (dynein) motor have repeatedly been implicated in neurological diseases (1, 2, 5, 7–16). Dynein is a 1.4 MDa complex that is responsible for the vast majority of cargo transport towards the minus ends of microtubules (17–19). The complex consists of two copies each of the 520 kDa heavy chain (DYNC1H1), an intermediate chain (either the DYNC1I1 or DYNC1I2 isoform), a light intermediate chain (either the DYNC1LI1 or DYNC1LI2 isoform), and three different light chains (DYNLT1 (Tctex1), DYNLL1 (LC8) and DYNLRB1 (Robl)) (17–19) (Figure 1A and B). The heavy chain contains a motor domain, which translocates the protein complex towards the minus ends of microtubules using the energy derived from ATP hydrolysis (19, 20). Key elements within the motor domain include a ring formed by six AAA+ domains, of which AAA1 is the major ATP hydrolysis site, a microtubule-binding domain (MTBD), and an antiparallel coiled-coil stalk that mediates communication between the ring and the MTBD. In addition, there is a buttress domain that emerges from AAA5 of the ring and provides support to the base of the stalk, a linker that is remodelled to produce the powerstroke, and a C-terminal domain that regulates force production and processivity (21). The heavy chain also contains the tail domain, which mediates dimerisation and provides a platform for recruiting the other chains. The accessory chains are important for linking the motor to cargoes, either through direct interactions or by binding to an intermediary cargo adaptor (22, 23).

At the time of writing, more than 30 heterozygous missense mutations in the *DYNC1H1* gene have been identified in patients diagnosed with spinal muscular atrophy with lower extremity predominance (SMALED (OMIM: 158600)) or malformations of cortical development (MCD (OMIM: 614563)) (1, 2, 5, 7–16). SMALED is characterised by muscle weakness in the legs, which is caused by congenital or early childhood-onset loss of spinal cord motor neurons. MCD arises from defective neuronal proliferation or migration during development of the cerebral cortex, and is often associated with severe intellectual disability and epilepsy. Whereas some mutations segregate with disease in familial cases (2, 7, 9–11, 16, 24), others have arisen *de novo* (2, 8, 10, 12–15) (Supplementary Table 1). For the vast majority of mutations, direct evidence of an effect on dynein function has not been provided and thus it has not been conclusively shown that they are pathogenic.

**Figure 1.**
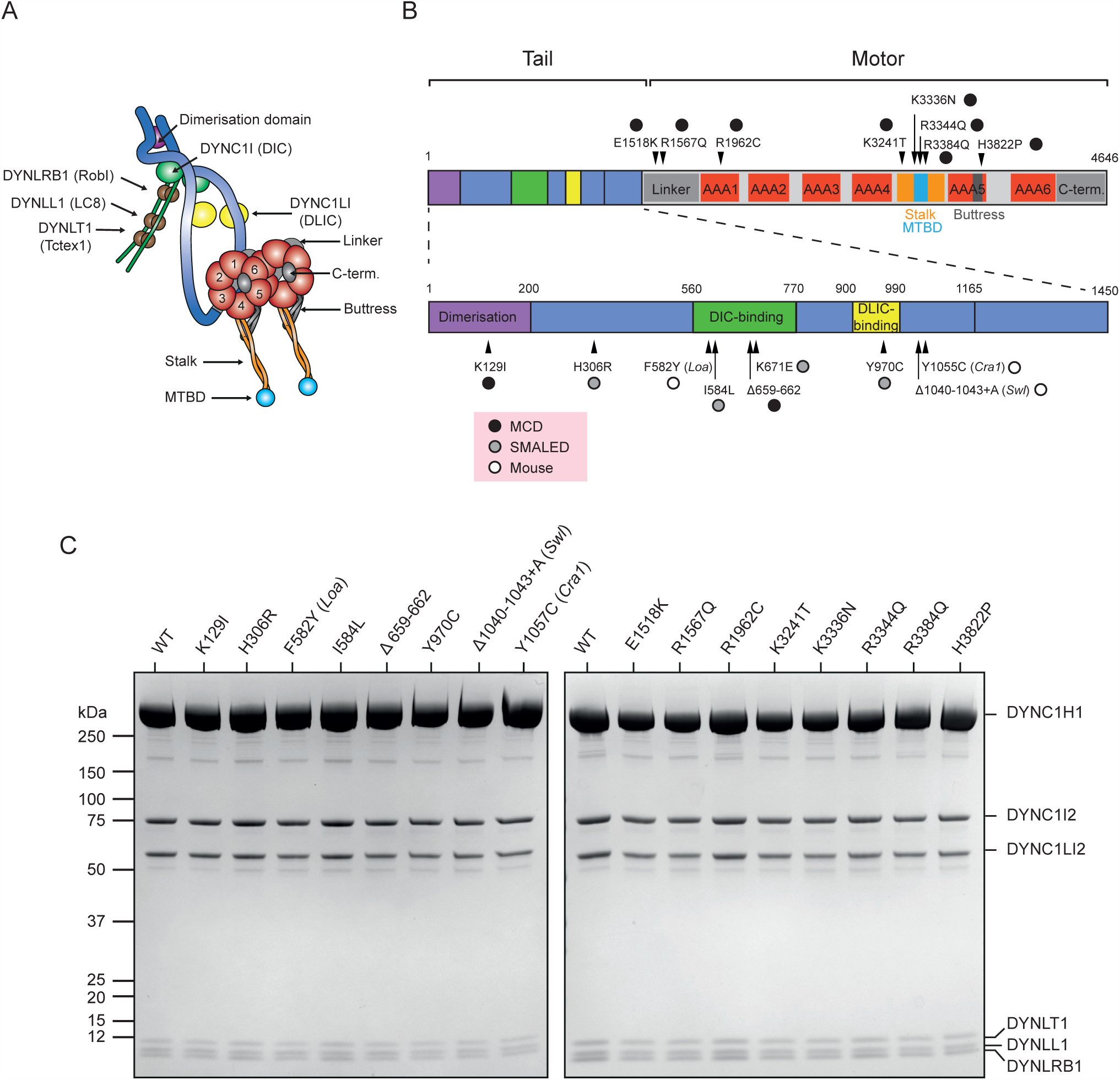
Positions of DYNC1H1 mutations and accessory chain composition of purified mutant dynein complexes. (A) Architecture of the full dynein complex. The linker domain is behind the AAA+ ring in this view. C-term: C-terminal domain. (B) Positions in the DYNC1H1 polypeptide of the human and mouse mutations characterised in this study. Mutations discovered in mouse are numbered according to the position of the equivalent residues in the human DYNC1H1 sequence. Note that H306R has been associated with axonal Charcot-Marie-Tooth disease type 2O in one pedigree (7). (C) Coomassie-stained denaturing gels of human recombinant dynein complexes purified from insect cells. The same amount of total protein was loaded per well. None of the DYNC1H1 mutations resulted in an overt change in accessory chain composition of purified dynein complexes compared to the wild-type (WT). This conclusion was confirmed with dynein samples expressed and purified from an independent construct. Note that K671E DYNC1H1 could not be purified from insect cells in three independent experiments using two different expression constructs.

The links between *DYNC1H1* mutations and neurological disease are strengthened further by the discovery of three heterozygous mutations that cause overlapping neuromuscular and sensory deficits in adult mice (1, 25–28). The *DYNC1H1* mutant strains were identified in genetic screens due to hind limb clenching and an abnormal gait (1), suggesting parallels with the SMALED phenotype in humans. Heterozygosity for a null mutation in *DYNC1H1* does not cause overt phenotypes in mice (29). This observation, together with the failure to detect nonsense, frameshift or deletion alleles of *DYNC1H1* in humans with SMALED or MCD, indicate that disease-associated mutations have a dominant-negative or dominant gain-of-function effect. However, with the exception of the mouse *Legs at odd angles (Loa)* mutation (27, 30–33), the effects of human and mouse mutations on the motility of the dynein complex have not been investigated in detail.

*In vitro* motility assays with purified motors have provided many valuable insights into microtubule-based transport mechanisms. By allowing single protein complexes to be studied in a defined context, these approaches allow molecular mechanisms to be investigated in a way that is very challenging in a cell. Here, we assay the effects of 14 *DYNC1H1* patient mutations and the three mouse mutations on the expression and motility of dynein complexes *in vitro*. These experiments take advantage of two recent advances. First, we exploit a recombinant expression system for the entire human dynein complex (34). This system, which involves the production of the motor complex from a single baculovirus in insect cells, greatly facilitates the purification of dynein complexes with defined mutations. Second, we build on the recent discovery of a way to study robust minus end-directed movement of individual dynein complexes *in vitro*. Whereas individual mammalian dyneins *in vitro* very rarely exhibit processive behaviour (the ability to take repeated steps along microtubules) (34–37), dynein complexes bound to the accessory complex dynactin and a cargo adaptor frequently move processively over very long distances (34, 37–39). Once bound to dynactin and a cargo adaptor, processive movements of dynein are faster and the motor has a greater force output (37, 40).

The best characterised of these activating cargo adaptors is Bicaudal-D2 (BICD2) (23, 41). Studies in rodents have shown that this protein plays an important role in dynein-based cargo transport in neurons (42–44). Consistent with these findings, missense mutations in the human *BICD2N* gene are also associated with SMALED (45–47) and cortical malformations (48, 49). The protein uses an N-terminal coiled-coil domain (BICD2N) to bind dynein and dynactin (50–52) and a C-terminal coiled coil (BICD2C) to bind cargoes. BICD2’s cargoes include nuclei that migrate apically in cortical neurons (43, 44, 53) and Golgi-derived vesicles (54). Structural electron microscopy has revealed how BICD2N bridges the interaction between dynein and dynactin (55). The presence of coiled-coil domains in other activating cargo adaptors suggests that they interact with dynein and dynactin through a related mechanism (37–39).

The results of our experiments provide insight into the relationship between *DYNC1H1* mutations and neurological defects. Our finding that all analysed *DYNC1H1* mutations compromise either dynein expression or motility provides functional evidence for their causative role in disease. We show that the vast majority of mutations inhibit the processive behaviour of dynein-dynactin-BICD2N complexes. Mutations with the strongest effects on dynein motility are associated with MCD in humans. Collectively, our results suggest that defective movement of cargo-motor complexes contributes to the neurological phenotypes associated with many *DYNC1H1* mutations. The functional effects of the mutations also reveal that many regions of the DYNC1H1 polypeptide regulate processivity of dynein-dynactin-cargo adaptor complexes. Several of these effects cannot be explained by our current understanding of dynein mechanism, and thus pave the way for further mechanistic studies.

## RESULTS

### Expression and purification of mutant dynein complexes

We focused our study on 17 disease-associated mutations in the *DYNC1H1* gene that were documented at the onset of the project (1) (Figure 1B; Supplementary Tables 1 and 2). This set includes 14 human mutations that are found in individuals diagnosed with MCD or SMALED and the three mouse mutations, *Cramping 1 (Cra1), Loa* and *Sprawling (Swl)*. 15 of these mutations result to single amino acid substitutions and two (Δ659-662 and Swl) produce small deletions in the DYNC1H1 polypeptide (Figure 1A). The affected amino acids are located throughout the tail and motor domains of DYNC1H1 (Figure 1B). More information on the phenotypes associated with these mutations in humans and mice is provided in Supplementary Tables 1 and 2.

We set out to produce 17 different recombinant human dynein complexes, each containing a unique disease-associated mutation in both copies of DYNC1H1. Mutations were introduced individually by site-directed mutagenesis into a plasmid containing *DYNC1H1* sequences. Mutant *DYNC1H1* genes were then transferred into a plasmid encoding the dynein accessory chains (see Materials and Methods for details of isoforms used), and the resultant construct transposed into a baculoviral genome (34). A tag introduced on the N-terminus of DYNC1H1 allows affinity purification of mutant dynein complexes from baculovirus-infected insect cells and covalent labelling with bright fluorophores (34).

16 of the 17 mutant dynein complexes could be purified from insect cells by size exclusion chromatography of samples released from the affinity matrix. The exception was the dynein complex containing the K671E mutation, which is located in the region of the tail proposed to associate with the intermediate chain (Figure 1A). In three independent experiments, K671E DYNC1H1 could not be detected during the size exclusion chromatography process, indicating a defect in protein expression or stability.

It has been reported that native dynein complexes isolated from I584L and *Loa* mutant mammalian cells have an altered composition compared to those isolated from wild-type cells (11, 56), although an independent study of native *Loa* dynein found no effect on complex assembly (32). We used our purified protein complexes to determine if any of the mutations prevent the interaction of DYNC1H1 with one or more of the dynein accessory chains in our expression system. The samples were run on denaturing gels and stained with Coomassie Blue. None of the mutations led to an overt reduction in the amount of the accessory chains co-purified with DYNC1H1 (Figure 1C). Collectively, our experiments demonstrate that the vast majority of the DYNC1H1 mutations do not prevent expression of the polypeptide or its association with the other dynein chains. The K671E mutation, however, strongly affects the expression or stability of DYNC1H1.

### R1962C and H3822P strongly inhibit microtubule gliding by surface immobilised human dynein complexes

We next used a microtubule gliding assay (35, 57, 58) to assess the consequences of the DYNC1H1 mutations on the core mechanical properties of the dynein complex. This assay involves immobilising dyneins on a glass surface and incubated them with free microtubules in the presence of ATP (Figure 2A). The collective minus end-directed activity of teams of dyneins results in translocation of microtubules along the surface of the imaging chamber (35, 57, 58).

**Figure 2.**
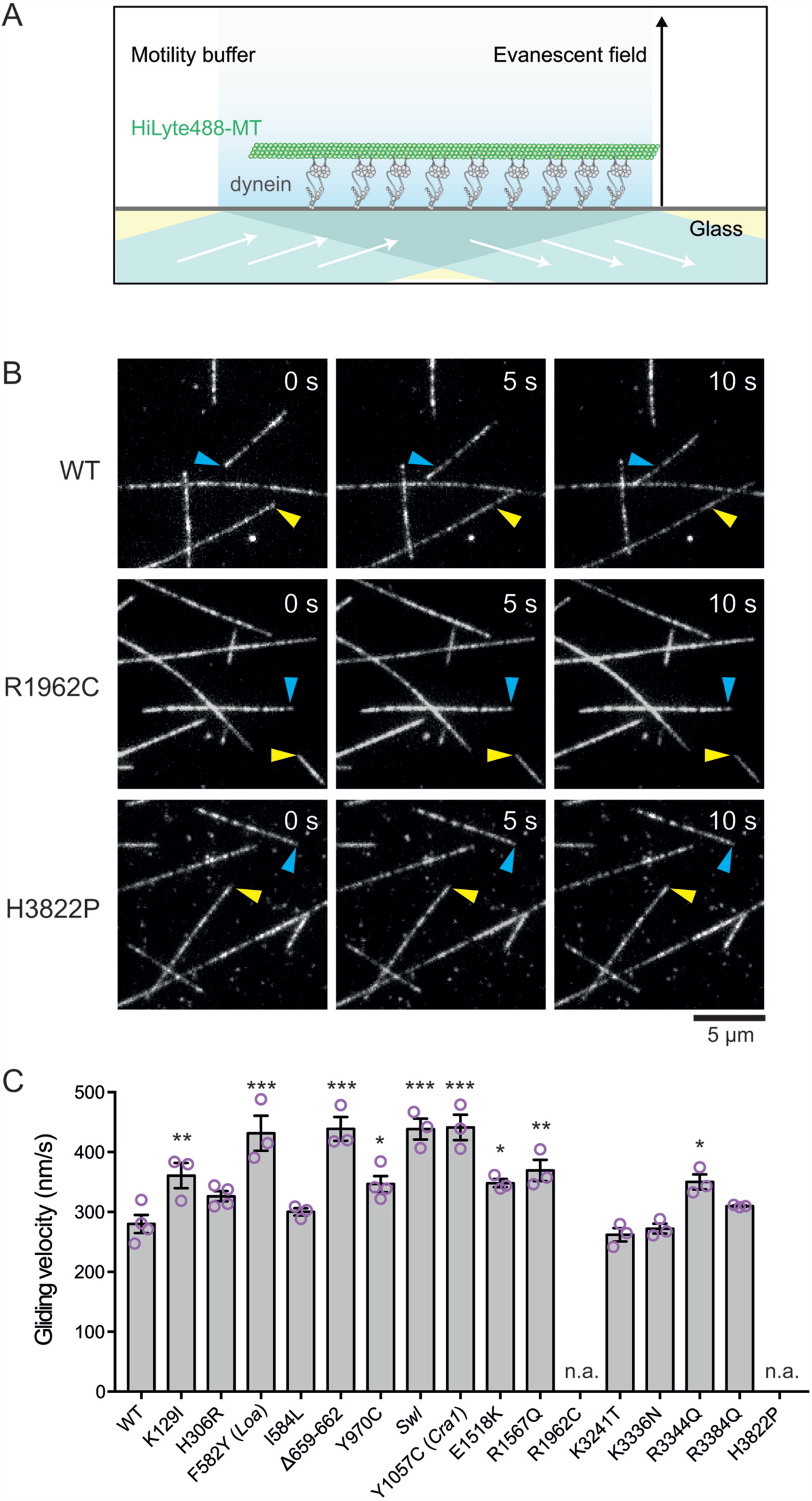
Effects of DYNC1H1 mutations on microtubule gliding by ensembles of human dynein complexes. (A) Diagram of the microtubule gliding assay. Dyneins are absorbed non-specifically onto the glass surface. MT: microtubule. (B) Stills from image series in Supplementary Movie 1 showing microtubules being translocated by wild-type (WT) dynein but not by dynein containing R1962C or H3822P DYNC1H1. In each condition, the starting position of two microtubule ends are labelled with arrows. (C) Quantification of gliding velocities in the presence of WT and mutant dynein complexes. Each magenta circle represents the mean value for an individual chamber (at least 30 microtubules analysed per chamber). Grey bars show means of the individual chamber values for each condition, with error bars representing S.E.M. Statistical significance, compared to WT, was evaluated with a one-way ANOVA with Sidak’s multiple comparison test (***, p < 0.001; **, < p <0.01; *, p < 0.05; n.a.: not applicable because to the absence (R1962C) or extreme rarity (H3822P) of microtubule gliding).

Fluorescent microtubules were used in our assays, which can be readily visualised by total internal reflection fluorescence (TIRF) microscopy. In the absence of dynein, microtubules did not adhere to the glass surface (Supplementary Movie 1). Preadsorption of the glass with wild-type dynein or any of the 16 obtainable mutant complexes resulted in persistent association of microtubules with the surface (e.g. Supplementary Movie 1). Thus, none of the mutations abolish the microtubule-binding activity of dynein. As expected, wild-type dyneins induced robust gliding of surface-associated microtubules (Figure 2B and C; Supplementary Movie 1). 14 of the 16 mutant dynein complexes also translocated microtubules robustly, doing so with velocities that were at least as high as those observed for wild-type dynein (Figure 2C).

In contrast, two mutants – R1962C and H3822P, which are in AAA1 and AAA5, respectively – exhibited a very strong defect in microtubule gliding. R1962C dyneins did not produce any movement of microtubules along the glass surface (Figure 2B; Supplementary Movie 1), whereas 94% of microtubules (117 out of a total of 124 analysed) were static when H3822P dynein was used (Figure 2B; Supplementary Movie 1). Those microtubules that were moved by H3822P dynein had a strongly reduced velocity compared to wild-type dynein (mean ± S.E.M.: H3822P, 30 ± 6 nm/s (N = 7 microtubules); wild-type, 280 ± 15 nm/s (N = 173 microtubules)).

To test the validity of these results, we produced each of the 16 mutant dyneins from an independent baculovirus construct. The strong inhibitory effects of the R1962C and H3822P mutations on microtubule gliding were confirmed when these new protein preparations were used, as was the ability of the other 14 mutations to support robust gliding (Supplementary Figure 1A and B). Interestingly, several of the dynein mutations in the tail or linker domains – K129I, F582Y (*Loa*), Δ659−662, Δ1040−1043+A (*Swl*), Y1057C (*Cra*), E1518K and R1567C – resulted in a substantial increase in the mean microtubule gliding velocity compared to the wild-type in both experimental series (Figure 2C; Supplementary Figure 1A and B). There was, however, no correlation between the effects of human mutations on microtubule gliding and their association with MCD or SMALED. We conclude from these experiments that the vast majority of dynein mutations do not inhibit motor activity of the human dynein complex in ensemble microtubule gliding assays. R1962C and H3822P, however, strongly inhibit dynein activity in this context.

### K129I, K3336N, R3344Q and R3384Q compromise activation of processive movement in the presence of dynactin and BICD2N

We next investigated the effects of the DYNC1H1 mutations on the processive motion of single dynein complexes. As described in the Introduction, processive motion of single mammalian dyneins is rarely observed in the absence of dynactin and a cargo adaptor (34–37). We therefore assayed the motility of the mutant dynein complexes in the presence of both the native dynactin complex, which was purified from pig brain, and recombinant BICD2N (amino acids 1-400) (see Supplementary Figure 2 for data on typical purity of these protein samples). To allow visualisation of protein complexes, dyneins were labelled with Tetramethylrhodamine (TMR) and BICD2N was labelled with Alexa647 (see Materials and Methods). Wild-type or mutant TMR-dyneins were incubated with dynactin and Alexa647-BICD2N, and protein mixes injected into imaging chambers containing surface-immobilised fluorescent microtubules. TIRF microscopy was then used to capture microtubule-associated TMR-dynein and Alexa647-BICD2N signals (Figure 3A).

**Figure 3.**
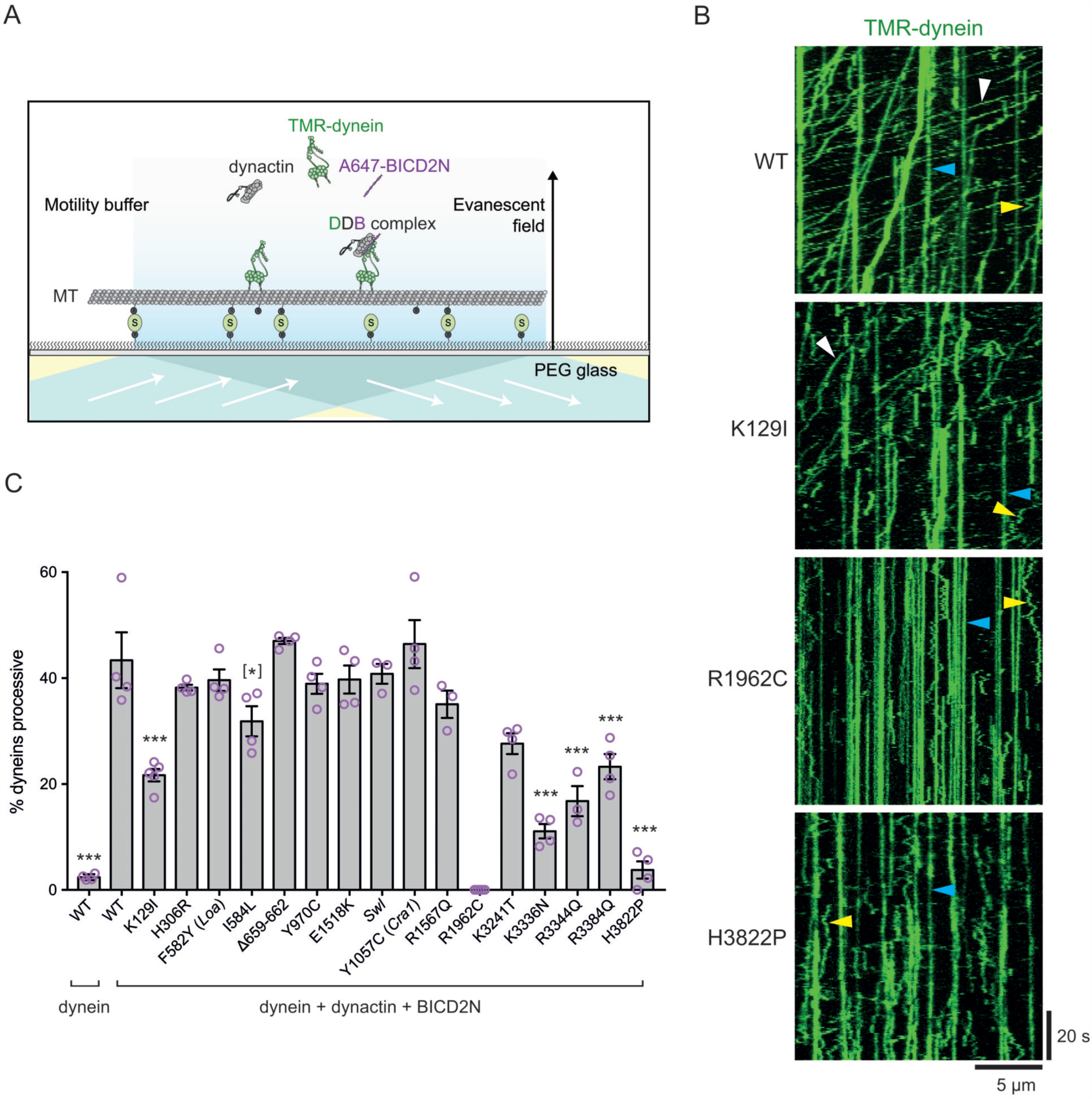
Effects of DYNC1H1 mutations on processive movement of individual dynein complexes in the presence of
dynactin and BICD2N. (A) Diagram of the assay used to monitor processive movement of dynein. Microtubules (MT) are immobilised on streptavidin (S)-coated glass by virtue of the biotin present on a fraction of tubulins. (B) Kymographs (time-distance plots) showing examples of the effect of DYNC1H1 mutations on processive movement of dynein (green) in the presence of dynactin and BICD2N. WT: wild-type. White, yellow and blue arrowheads indicate examples of processive, diffusive and static behaviour, respectively. (C) Quantification of the effects of DYNC1H1 mutations on the percentage of microtubule-associated dyneins that move processively in the presence of dynactin and BICD2N. Each magenta circle represents the mean value for an individual chamber (at least 70 dynein complexes analysed per chamber). Throughout the study, each chamber used an independent assembly mix of dynein, dynactin and BICD2N. Grey bars show means of the individual chamber values for each condition, with error bars representing S.E.M. Statistical significance, compared to WT dynein in the presence of dynactin and BICD2N, was evaluated with a one-way ANOVA with Sidak’s multiple comparison test (***, p < 0.001; *, p < 0.05; [*], p<0.05 but a significant difference was not reproduced in a second experimental series (Supplementary Figure 3)).

We first analysed the TMR signals in order to assess the behaviour of the total dynein population on microtubules. In the presence of dynactin and BICD2N, ˜40% of microtubule-associated dynein complexes containing wild-type DYNC1H1 underwent processive movement (Figure 3B and C). Less than 3% of the wild-type dyneins on microtubules exhibited processive motion when dynactin and BICD2N were omitted (Figure 3C). These values were similar to those observed in our previous study of wild-type dynein in the presence and absence of dynactin and BICD2N (34).

As expected from their effects on microtubule gliding by dynein, the R1962C and H3822P mutations strongly inhibited processive motion of single dynein complexes in the presence of dynactin and BICD2N (Figure 3B and C). No processive complexes were observed for R1962C dynein (Figure 3C), whereas an average of 5% of microtubule-associated H3822P complexes exhibited processive movement (Figure 3C). Thus, although both R1962C and H3822P mutations had very strong effects on the behaviour of single dynein complexes in the presence of dynactin and BICD2N, only the R1962C mutation completely abolished motility. Four other DYNC1H1 mutations – K129I, K3336N, R3344Q and R3384Q – resulted in only 10–20% of TMR-dynein complexes undergoing processive motion in the presence of dynactin and BICD2N, which equates to an ˜ 2 to 4-fold reduction compared to the wild-type (Figure 3B and C). K129I is found in the N-terminal region of the DYNC1H1 tail and K3336N, R3344Q and R3384Q are found in the MTBD (Figure 1B). The tail mutation I584L resulted in a smaller, but statistically significant, reduction in the frequency of processive motion in the presence of dynactin and BICD2N compared to the wild-type (Figure 3C). The remaining mutations had no effect on the frequency of processive motion (Figure 3C). We confirmed that the K129I, R1962C, K3336N, R3344Q, R3384Q and H3822P mutations reduced the frequency of processive movements in another experimental series, which used independent preparations of the 16 mutant dynein complexes, dynactin and BICD2N (Supplementary Figure 3A and B). However, the subtle effect of the I584L mutation observed previously was not reproduced in this series (Supplementary Figure 3B).

In summary, these experiments demonstrate that several DYNC1H1 mutations reduce the incidence of processive behaviour of single dynein complexes that is induced by dynactin and BICD2N. The strongest effects were observed for R1962C and H3822P, which is consistent with the strong inhibitory effects of these mutations on microtubule gliding. K129I, K3336N, R3344Q and R3384Q, which did not compromise dynein activity in the gliding assay, consistently led to a strong reduction in the incidence of processive movement of dynein complexes.

### Most DYNC1H1 mutations do not inhibit binding of dynein to dynactin and BICD2N

A decreased incidence of processive movement by some mutant dyneins could be caused by reduced association with dynactin and BICD2N, or by compromised motility of dynein-dynactin-BICD2N complexes once they are assembled. To distinguish between these possibilities, we determined the percentage of microtubule-associated dynein complexes containing each DYNC1H1 mutation that associated with dynactin and BICD2N. This was achieved by overlaying the Alexa647-BICD2N signals on the TMR-dynein signals (Figure 4A; Supplementary Figures 4 and 5) and scoring co-localisation. As BICD2N can only bind dynein in the presence of dynactin (34, 37, 52), dual coloured puncta must represent dynein-dynactin-BICD2N complexes. Consistent with this notion, whereas ˜75% of the total population of wild-type TMR-dyneins had an Alexa647-BICD2N signal when dynactin was included in the assay (Figure 4B), no co-localisation was observed when dynactin was excluded (Supplementary Figure 6). As expected, in the presence of dynactin, almost all processive wild-type dynein complexes had a detectable signal from BICD2N (Figure 4A and Supplementary Figure 4). However, not all wildtype dyneins with a BICD2N signal moved processively (Figure 4A and C), in agreement with previous observations that not all dynein-dynactin-BICD2N complexes are active (34, 37).

Co-localisation analysis for mutant complexes revealed that only two DYNC1H1 mutations – K3241T and K3336N – significantly reduced the degree of co-localisation of dynein and BICD2N on microtubules (Figure 4B). The magnitude of these effects was, however, rather small: approximately 60% of K3241T dyneins associated with dynactin and BICD2N, with this value decreasing to 50% for K3336N (Figure 4B). We conclude that the majority of dynein mutations do not alter the ability of dynein to associate with dynactin and BICD2N *in vitro*.

**Figure 4.**
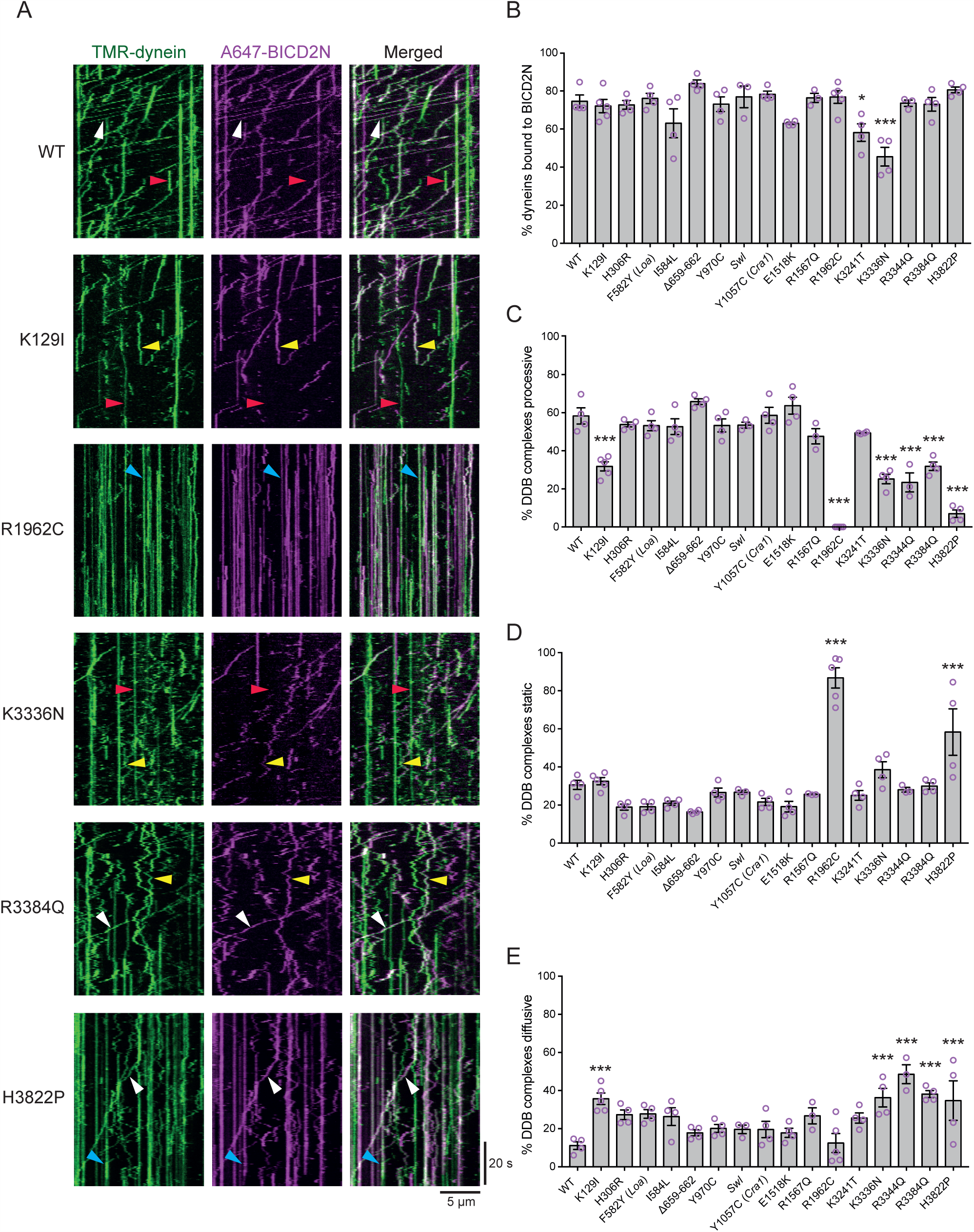
Effects of DYNC1H1 mutations on the assembly and processivity of dynein-dynactin-BICD2N complexes. (A) Kymographs showing examples of frequent co-localisation of mutant TMR-dynein complexes (green) with Alexa647-BICD2N (magenta) on microtubules in the presence of dynactin (see Supplementary Figures 4 and 5 for examples for other mutant complexes). Co-localisation indicates formation of a dynein-dynactin-BICD2N complex. White, blue and yellow arrowheads indicate examples of co-localisation of Alexa647-BICD2N with TMR-dynein exhibiting processive, static and diffusive behaviour, respectively. Red arrowheads indicate examples of TMR-dynein complexes that do not have an Alexa647-BICD2N signal. (B) Quantification of the percentage of all microtubule-associated dyneins that co-localises with BICD2N in the presence of dynactin. (C–E) Quantification of the percentage of microtubule-associated dynein-dynactin-BICD2N (DDB) complexes that exhibits processive (C), static (D) and diffusive (E) behaviour. In B–E, each magenta circle represents the mean value for an individual chamber (at least 70 dynein complexes (B) and 50 dynein-dynactin-BICD2N (C–E) complexes analysed per chamber). Grey bars show means of the individual chamber values for each condition, with error bars representing S.E.M. Statistical significance, compared to WT, was evaluated with a one-way ANOVA with Sidak’s multiple comparison test (***, p < 0.001; *, p < 0.05).

These data revealed that the K129I, R1962C, R3344Q, R3384Q and H3822P mutations do not inhibit the processive motion of dynein by interfering with dynein’s association with dynactin and BICD2N, but rather by compromising the motility of dynein-dynactin-BICD2N complexes. The effect of these mutations on the ability of dynein-dynactin-BICD2N complexes to initiate processive movement was confirmed directly by analysing the motility of those TMR-dyneins with an Alexa647-BICD2N signal (Figure 4C). Interestingly, the mutation that exhibited the strongest reduction in the co-localisation of dynein and BICD2N – K3336N – also significantly reduced the frequency of processive motion of those dynein-dynactin-BICD2N complexes that were formed (Figure 4C).

We and others have noted that non-processive dynein complexes can be statically bound to microtubules or undergo short-range diffusion along the lattice (34–37). This latter behaviour is likely to reflect a weak interaction of the motor complex with the microtubule (36). The reduction in processive motion of dynein-dynactin-BICD2N complexes containing the K129I, K3336N, R3344Q and R3384Q mutations was accompanied by a significant increase in the frequency of diffusive behaviour (Figure 4C–E). In contrast, the vast majority of the dynein-dynactin-BICD2N complexes containing the R1962C mutation in AAA1 exhibited static binding to microtubules (Figure 4C–E; Supplementary Figure 3A). The H3822P mutation led to an increase in the incidence of both static and diffusive events (Figure 4C–E). Collectively, these data indicate that several of the DYNC1H1 mutations affect the probability of processive, diffusive and static behaviour of dynein-dynactin-BICD2N complexes on microtubules.

### The vast majority of dynein mutations reduce the travel distance of dynein-dynactin-BICD2N complexes

We next examined the effects of the DYNC1H1 mutations on the run length and velocity of those dynein-dynactin-BICD2N complexes that exhibited processive motion. R1962C dyneins were excluded from this analysis as they displayed no processive movements. Because processive movements were very rare for H3822P dyneins, we had to pool run length and velocity values across chambers for this mutant (Supplementary Figure 7) rather than performing chamber-by-chamber analysis.

Operationally, run length was defined as the distance a processive complex travelled before pausing or detaching from the microtubule. Strikingly, all 15 mutations analysed resulted in an approximate two-fold reduction in the mean run length of dynein-dynactin-BICD2N complexes (Figure 5A, B; Supplementary Figure 7 and 8A). The ability of DYNC1H1 mutations to reduce run lengths was confirmed by analysis of the *in vitro* motility assays performed with independent preparations of dynein complexes, dynactin and BICD2N (Supplementary Figure 9A–C). Thus, all DYNC1H1 disease mutations decrease the distance of processive travel of dynein-dynactin-BICD2N complexes on microtubules.

**Figure 5.**
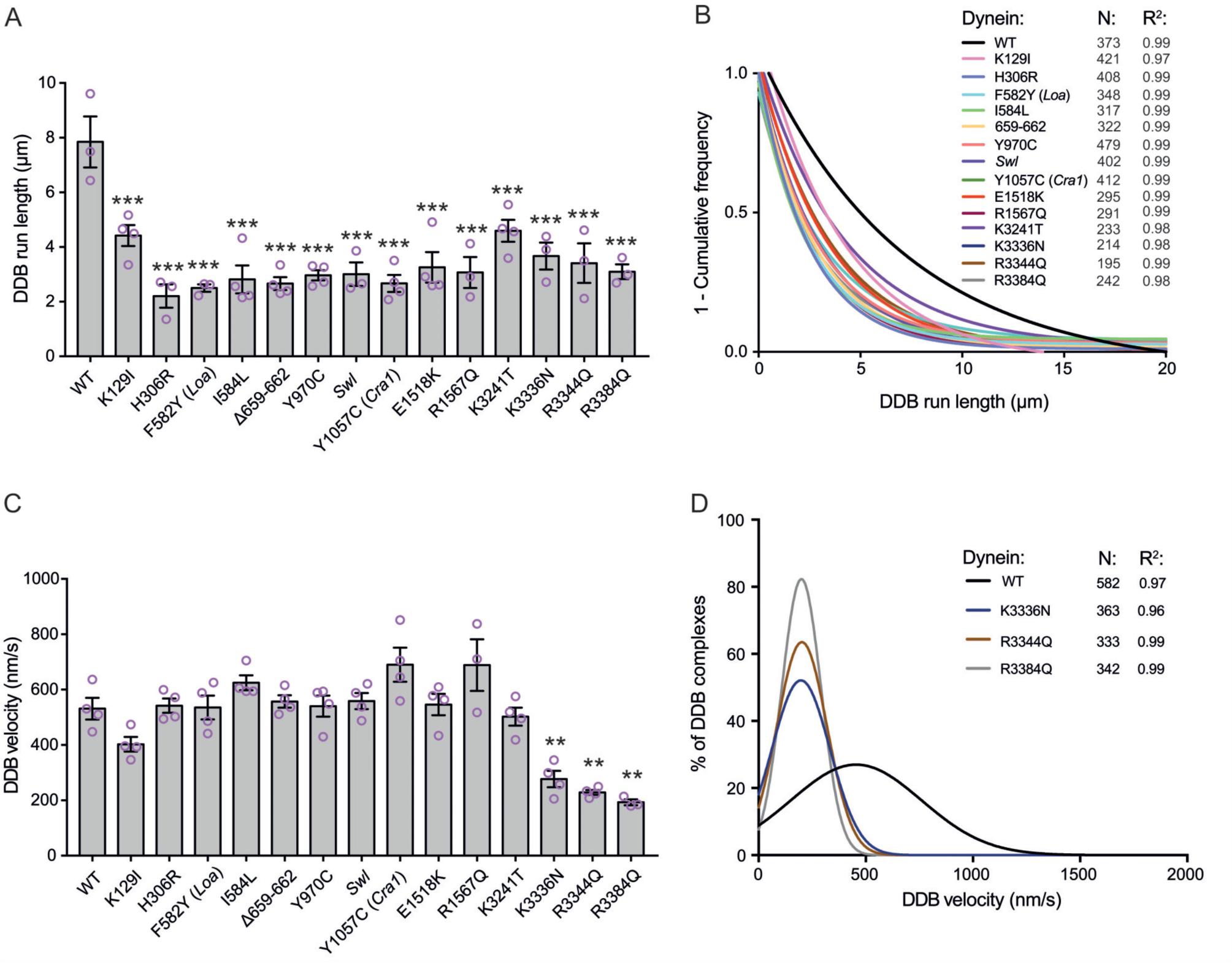
Effects of DYNC1H1 mutations on the run length and velocity of processive dynein-dynactin-BICD2N complexes. Quantification of the effects of DYNC1H1 mutations on the run length (A, B) and velocity (C, D) of processive dynein-dynactin-BICD2N (DDB) complexes. In A and C, each magenta circle represents the mean value for an individual chamber (at least 50 runs or velocity segments were analysed per chamber). Grey bars show means of the individual chamber values for each condition, with error bars representing S.E.M (WT: wild-type). Statistical significance, compared to WT, was evaluated with a one-way ANOVA with Sidak’s multiple comparison test (***, p < 0.001; **, < p <0.01). B and D show the distributions of run lengths and velocities summed across all chambers for those DYNC1H1 mutations with a sizeable effect. B shows curves fitted to a ‘1 minus cumulative frequency’ plot, after McKenney et al. (37). D shows curves fitted to the raw data. N: total number of runs (B) or velocity segments (D). R^2^: goodness of fit. See Supplementary Figure 8 for examples of raw run length and velocity distributions. Due to the strong reduction in the frequency of processive motion, there were insufficient data points to plot the run lengths and velocities for H3822P dyneins on a per chamber basis. The pooled values across all chambers were therefore analysed separately (Supplementary Figure 7).

We next analysed the effects of the mutations on the velocity of processive dynein-dynactin-BICD2N complexes. The K3336N, R3344Q, R3384Q and H3822P mutations led to an approximately two-fold decrease in mean velocity of dynein-dynactin-BICD2N complexes (Figure 5C and D; Supplementary Figures 7, 8B, 9D and E). These mutations also consistently reduced the frequency of processive transport events by dynein-dynactin-BICD2N complexes (Figure 3C). Thus, K3336N, R3344Q, R3384Q and H3822P affect multiple aspects of dynein motility. The other 11 mutations analysed did not consistently affect the velocity of processive dynein-dynactin-BICD2N complexes (Figure 5C and Supplementary Figure 9D). Thus, the ability of several tail and linker mutations to increase the velocity of microtubule gliding by teams of immobilised dyneins were not accompanied by an increase in the speed at which dynein-dynactin-BICD2N complexes run along microtubules. Our finding that the *Loa* mutation reduces run lengths but not velocities of a dynein-dynactin-cargo adaptor complex are consistent with results of high spatiotemporal imaging of lysosomes in *Loa* mutant mouse axons (32).

Overall, we conclude from our analyses that the vast majority of disease-associated DYNC1H1 mutations impair dynein function not by inhibiting binding to processivity activators, but rather through restriction of movement of the intact transport complex.

## DISCUSSION

### Summary

We have used a recently developed insect cell expression system and *in vitro* motility assays to characterise the effects on dynein motility of a large series of mutations in DYNC1H1 that are associated with neurological disease in human and mouse. The motility assays included an assessment of long-distance motility of individual dynein complexes, which was made possible by the discovery that processive movement of mammalian dynein *in vitro* can be activated by its binding to the dynactin complex and a cargo adaptor, such as BICD2 (34, 37). Our results are summarised in Figure 6. We identify two mutations – R1962C and H3822P – that severely compromise the core mechanochemical properties of human dynein as judged by their strong effects on microtubule gliding by ensembles of dyneins, as well as on processive movement of individual motor complexes in the presence of dynactin and BICD2N. Four mutations – K129I, K3336N, R3344Q and R3384Q – do not inhibit microtubule gliding by human dynein but consistently reduce the frequency of processive movement of the motor complex in the presence of dynactin and BICD2N. These mutations also strongly reduce the travel distance of processive events, with K3336N, R3344Q and R3384Q additionally leading to a significant slowing of these movements. A further ten mutations, including all three that were discovered in the mouse, had a more restricted effect on the behaviour of human dynein in our assays. The predominant effect of these mutations was a substantial decrease in the length of processive transport events in the presence of dynactin and BICD2N. Overall, the mutations studied had little effect on the ability of dynein to associate with dynactin and BICD2N, suggesting that they result in BICD2’s cargoes being linked to dynein complexes with defective motility.

**Figure 6.**
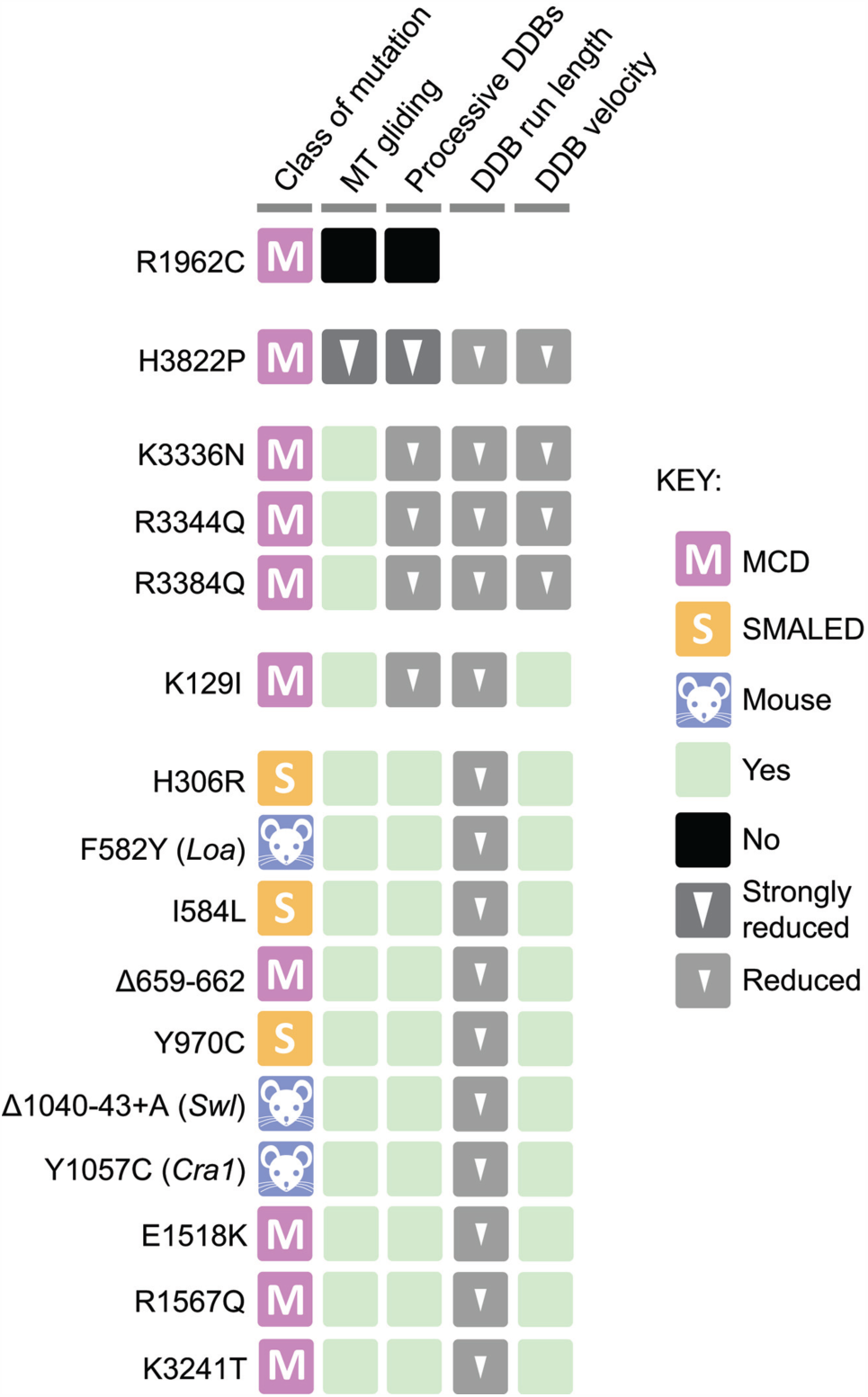
Summary of the consequences of DYNC1H1 mutations on dynein motility. Mutations are clustered based on similarities of their effects on purified dynein complexes, with the severity of each cluster decreasing from top to bottom. Within each cluster, mutations are ordered in an N–C terminal direction. MT: microtubule; DDB: dynein-dynactin-BICD2N. Note that K671E DYNC1H1 could not be purified.

### Mechanistic effects of disease-associated DYNC1H1 mutations

Although structural and single molecule studies have provided remarkable insights into dynein mechanism (18–20), the dynamic changes within the motor complex that generate movement of cargoes along microtubules are only partially understood. Our analysis of disease-associated mutations highlight several positions of the ˜4600 amino acid-long DYNC1H1 that are important for efficient motility of a dynein-dynactin-cargo adaptor complex.

Eight of the mutations we studied are located in the motor domain (Figure 7A). A high-resolution structure of the motor domain of mammalian DYNC1H1 has not been reported. However, homology modelling of the motor domain based on high-resolution structures from other dyneins (59, 60) or a pseudo-atomic model of the MTBD from mouse DYNC1H1 (61) offer an explanation for how several of the mutations affect motor function. The R1962C mutation is located within the large domain of AAA1 (AAA1L) and so affects the main ATPase site in dynein. In the presence of ADP there is a cleft between AAA1 and AAA2 and R1962 points into solution (Figure 7B). Upon ATP binding and hydrolysis the cleft closes and R1962 becomes part of a network of contacts between AAA1L and AAA2L (Figure 7C). The R1962C mutation would disrupt these contacts and is thus expected to destabilise the conformation dynein enters during ATP hydrolysis. This would account for the lack of R1962C dynein activity in the microtubule gliding assay, as well as in the context of individual dynein-dynactin-BICD2N complexes. Because closure of AAA1 and AAA2 also plays a key role in the structural rearrangements that lower the affinity of the MTBD for the microtubule (20), an inhibitory effect on this process could also explain why most R1962C dyneins are stably bound to a single site on the microtubule rather than engaged in lattice diffusion (Figure 4D and E).

**Figure 7.**
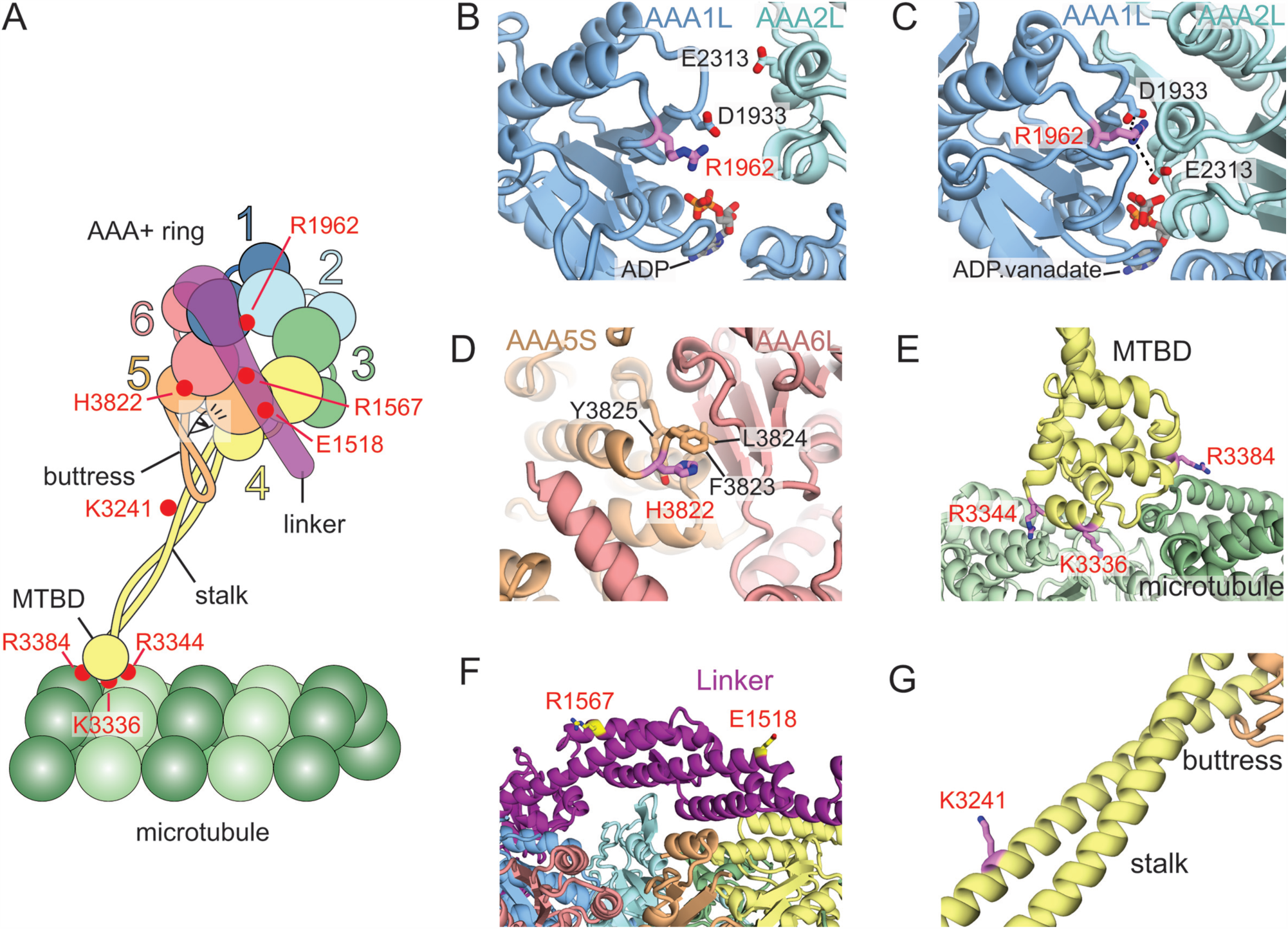
Structural analysis of mutated residues in the DYNC1H1 motor domain. (A) Cartoon of a microtubule-bound dynein motor domain, showing positions of DYNC1H1 mutations (red circles). Individual AAA+ domains are labelled with a coloured number. Each AAA+ domain is divided into a large subdomain and a small subdomain, which are indicated by circles of different sizes. MTBD: microtubule binding domain. Eye shows the viewpoint for panel F. (B–F) Structural analysis of the positions of mutated residues in the DYNC1H1 motor domain. Domains are coloured as in A; side chains of mutated residues are shown in magenta in B–E and G, and yellow in F. (B) R1962 in the AAA1 large domain (AAA1L) makes no contacts when dynein binds ADP in the AAA1 site. (C) R1962 makes a network of contacts (dashed lines) when the AAA1/AAA2 cleft closes in the presence of a nucleotide analogue (ADP.vanadate) that mimics the transition state of ATP hydrolysis. (D) H3822 is found in the AAA5 small domain (AAA5S). It packs against three residues that make contact with the AAA6 large domain (AAA6L). (E) K3336, R3344 and R3384 reside on the surface of the MTBD that binds the microtubule. (F) E1518 and R1567 are on the surface of the linker domain, pointing away from the AAA+ domains. (G) K3241 is on the surface of the stalk pointing into solution. Homology models are based on the structures of the motor domain of *Dictyostelium* dynein-1 bound to ADP (PDB:3VKG) (panels B, D, F and G), human cytoplasmic dynein-2 bound to ADP.vanadate (PDB:4RH7) (panel C) and a pseudo-atomic model from a 9 Å cryo-EM structure of the MTBD of mouse DYNC1H1 bound to microtubules (PDB:3J1T) (panel E).

H3822P has a similar affect to R1962C in the *in vitro* motility assays, with very little microtubule gliding activity or processive movement of dynein-dynactin-BICD2N complexes. H3822 is in the small domain of AAA5 (AAA5S) close to the interface with the large domain of AAA6 (AAA6L) (Figure 7D). AAA6L and AAA5S move together as a block during the ATP hydrolysis cycle of dynein (20). AAA5L also contains the buttress (Figure 7A), which couples these movements to changes in the conformation of the stalk and the affinity of the MTBD for microtubules. H3822P would disrupt a cluster of AAA5S residues (F3823, L3824 and Y3825) at the interface with AAA6L (Figure 7D). This mutation may therefore compromise dynein motility by interfering with the communication pathway that couples rearrangements in the AAA+ ring to changes in the MTBD.

As noted previously (2), three of the mutations we analysed – K3336N, R3344Q and R3384Q – lie at the interface between the MTBD and microtubules (Figure 7A and E) (61). All three mutations strongly reduce the frequency, travel distance and velocity of processive movements of individual dynein-dynactin-BICD2N complexes but have no effect on microtubule gliding by surface-immobilised dyneins. We found that these mutations significantly increase the frequency of lattice diffusion of dynein-dynactin-BICD2N complexes (Figure 4E), suggesting that they weaken the interaction between the MTBD and the microtubule. Consistent with this notion, K3336N and R3384Q reduce the co-sedimentation of the MTBD with microtubules in the context of a fusion protein with a heterologous coiled-coil domain (2). By interfering with the tight binding state of the MTBD, the K3336N, R3344Q and R3384Q mutations may disrupt the ability of the dynein heads to make repeated steps. This behaviour is essential for processive movement of individual complexes but is unlikely to be required for microtubule gliding by teams of surface-immobilised motors (35).

The three other disease-associated mutations in the motor domain – E1518K, R1567Q and H3241T – affect surface exposed residues. E1518K and R1567Q are located in the linker domain (Figure 7F), whereas K3241 is found within the stalk (Figure 7G). These mutations reduce the run length of dynein-dynactin-BICD2N complexes but do not significantly affect the frequency or velocity of processive movements. How these three mutations affect travel distance of dynein-dynactin-BICD2N complexes is currently not clear, but they could conceivably modulate the dynamic conformational changes exhibited by the linker and stalk during the ATPase cycle.

The remaining DYNC1H1 mutations we studied are in the tail domain, for which there is no high-resolution structural information. We anticipated that at least some of the mutations in the DYNC1H1 tail would affect dynein function by altering the interaction with the dynein accessory chains, dynactin or BICD2N. Our failure to detect changes in the ability of the mutant DYNC1H1 proteins to interact with these factors was therefore surprising. These mutations do, however, compromise processive movements of dynein-dynactin-BICD2N complexes (Figure 6).

There are precedents for mutations in the tail of DYNC1H1 having effects on the motor domain without affecting interactions with binding partners. Dyneins isolated from mice heterozygous for the *Loa* mutation have curtailed minus end-directed motion *in vitro* and evidence of disrupted co-ordination between motor heads, despite having a full complement of accessory chains (32). Because the study was performed in the absence of processivity activators it is, however, unclear how the findings relate to our observations of the effect of the *Loa* mutation on dynein-dynactin-BICD2N complexes. It has also be demonstrated that a mutation in the tail of DYNC1H1 in the filamentous fungus *Aspergillus nidulans* (in a residue equivalent to position 186 of the human protein) reduces the frequency and velocity of minus end-directed cargo transport *in vivo* without affecting the composition of the dynein complex or its association with dynactin (62). Our study reveals that the ability of the tail to regulate the behavior of the motor domain is not just restricted to the regions containing the *Loa* and *Aspergillus* mutations. We show that many sites along the DYNC1H1 tail are involved in regulating motor activity in the context of a defined dynein-dynactin-cargo adaptor complex. Our data also suggest that disruption of this regulation contributes to neurological diseases in humans. Remarkably, the tail mutation with the strongest effect on processive movements of the dynein-dynactin-BICD2N complex in our study – K129I – is the one furthest away from the motor domain. Although this mutation is found within the so-called ‘dimerisation domain’ (55), it does not disrupt dimerisation of the heavy chains (Supplementary Figure 10). Thus, this variant also compromises the activity of the motor domain in the context of the full dynein-dynactin-BICD2N complex. Our identification of a large number of mutations in the tail that affect processive movements of dynein-dynactin-cargo adaptor complexes will aid future efforts to understand the basis of the communication between the tail and motor domains. This work may also illuminate how some mutations in the tail increase the velocity of microtubule gliding by teams of isolated dyneins.

### Insights into the basis of DYNC1H1-associated neurological diseases

Many of the *DYNC1H1* mutations identified in patients have either arisen *de novo* or have been identified in small families where segregation studies cannot conclusively demonstrate pathogenicity. In these cases, the evidence for involvement in disease has also included the evolutionary conservation of the mutated residue and differences in the physicochemical properties of the substituted amino acid. However, functional assays are highly desirable when ascribing causality to rare sequence variants (63). We find that all 14 human *DYNC1H1* mutations tested, including several *de novo* mutations (Supplementary Table 1), compromise the expression or motility of the dynein complex in a defined *in vitro* system. These data provide functional evidence for their contribution to neurological disease. Interestingly, of the 16 human or mouse mutations that support production of the recombinant dynein complex, the six with the strongest effects on dynein motility are associated with MCD in humans (Figure 6). All three SMALED mutations that could be assayed in the context of the purified motor complex fell into the class with the weakest effects (Figure 6). These mutations are among the group that only compromised run lengths of processive dynein-dynactin-BICD2N complexes. These findings raise the possibility that MCD and SMALED are caused by different degrees of inhibition of dynein motility, rather than inactivation of distinct dynein-related processes.

Consistent with the notion that dynein-associated neurological diseases are on the same spectrum, at least some individuals with *DYNC1H1* mutations have combined defects in cortical development and spinal motor neurons (2, 7, 10, 11, 13). For example, some of the DYNC1H1 mutations we studied were found in individuals with both MCD and foot deformities consistent with axonal neuropathy ((2); Supplementary Table 1). Other mutations in our study were identified in SMALED patients noted to also have mild intellectual disability (7, 11) (Supplementary Table 1). However, four *de novo* MCD-associated mutations – Δ659-662, E1518K, R1567Q and K3241T – fell into the class that only exhibited reduced run lengths of dynein-dynactin-BICD2N complexes. The overt cortical development defects in the individuals with these mutations may therefore result from an interaction with other mutations present in their genomes or environmental factors. The notion that extrinsic factors influence the phenotypes associated with human DYNC1H1 mutations is supported by the observation that the H306R variant gives rise to different symptoms in different pedigrees (7, 9) (Supplementary Table 1). Alternatively, by impacting on interactions of DYNC1H1 with other binding partners, Δ659-662, E1518K, R1567Q and K3241T may exert stronger effects on dynein motility *in vivo* than were evident in our study.

A key finding of our work is that the vast majority of DYNC1H1 mutations do not inhibit the ability of dynein to associate with dynactin and BICD2N, but do compromise the motile properties of dynein-dynactin-BICD2N complexes. An ability of dyneins with compromised motility to bind BICD2, and potentially other cargo adaptors, offers a possible explanation for the dominant effects of *DYNC1H1* mutations in humans and mice. The mutant dyneins would be expected to provide a link between cargoes and microtubules. However, processive movements of these cargoes would be shorter, slower or less frequent, thus compromising cargo delivery within neurons. Provided levels of DYNC1H1 protein are not limiting, heterozygosity for a null mutation in *DYNC1H1* would not be expected to have the same effect. This offers a potential explanation for why mice that are heterozygous for a *DYNC1H1* null allele are asymptomatic and frameshift, nonsense and deletion alleles have not been recovered in humans with MCD or SMALED.

As noted previously (32), neurons with long axonal processes are likely to be particularly sensitive to reduced processivity of cargo-dynein complexes. This could account for the association of those *DYNC1H1* mutations that have relatively mild effects on dynein activity *in vitro* with SMALED, which predominantly affects spinal motor neurons in the legs. Mutations with stronger effects on dynein motility may additionally compromise cargo delivery in neurons with shorter processes, such as those in the developing cortex, which could contribute to MCD. In the future, it will important to test this hypothesis in animal models.

Not all DYNC1H1 mutations studied could have their functional effects ascribed to reduced motility of the dynein complex. We found one mutation – K671E – that consistently prevented the purification of DYNC1H1 from insect cells. This mutation is found within the region of the tail that associates with dynein’s intermediate chain. Depletion of the intermediate chain in cells causes a large reduction in the level of DYNC1H1, indicating that stability of the heavy chain is dependent on its ability to associate with the intermediate chain (64). Any ability of the K671E mutation to disrupt intermediate chain binding would therefore be expected to destabilise DYNC1H1. It is not clear why heterozygosity for this mutation is associated with a phenotype in humans – SMALED (11) – whereas heterozygosity for nonsense, frameshift or deletion alleles in *DYNC1H1* is not. However, because defects in protein homeostasis have frequently been linked to neurological diseases (65), it is tempting to speculate that the destabilising effect of the K671E mutation results in a misfolded or aggregrated intermediate that has a pathological effect in neurons.

### Perspective

Since this study was initiated, the number of *DYNC1H1* mutations that have been associated with human neurological disease has risen substantially. For example, *DYNC1H1* variants of unknown clinical significance were commonly found in patients undergoing genetic screening for peripheral neuropathy (66). The number of variants identified in this gene is set to increase further with the adoption of next generation sequencing in routine clinical practice. Our work describes a pipeline that can be used to evaluate if and how newly discovered mutations affect dynein function. The same strategy can also be used to study the functional effects of *BICD2* variants identified in patients. We anticipate that studying the effects of disease-associated variants *in vitro* will also provide novel mechanistic insights into how dynein-dynactin-cargo complexes move along microtubules in a non-pathological setting. Ultimately, *in vitro* studies of mutations, coupled to analysis in cellular systems, should inform therapeutic efforts to tackle dynein-associated neurological diseases.

## MATERIALS AND METHODS

### Cloning of *DYNC1H1* mutations

Mutations were introduced by site-directed mutagenesis into *pDyn1* (34), which contains human *DYNC1H1* sequences (accession number NM_001376.4) that have been codon optimised for Sf9 insect cell expression. The plasmid also codes for a ZZ-LTLT-SNAP domain at the N-terminus of DYNC1H1. The ZZ tag consists of two IgG-binding domains based on *Staphylococcus aureus* protein A, the LTLT region includes a cleavage site for the tobacco etch virus (TEV) protease, and the SNAP tag allows covalent labelling with fluorescent dyes. The same mutations were also introduced into a variant of *pDyn1* that has the SNAP tag replaced with a Halo tag, which can also be used for fluorescent protein labelling. The Halo-tagged versions of DYNC1H1 were used in the experiments documented in Supplementary Figures 1, 3 and 9, which were designed to test the effects of each of the *DYNC1H1* mutations on the motility of dynein expressed from an independent construct. Site-directed mutagenesis was performed with Phusion High-Fidelity or Q5 Hot Start High Fidelity DNA polymerases (New England Biolabs (NEB)). The presence of the desired mutation was confirmed by sequencing. Sequencing of the entire *DYNC1H1* open reading frame in each plasmid confirmed the absence of off-target mutations, with the exception of an E97D codon change that was inadvertently introduced into the Halo-tagged version of *DYNC1H1* H306R. Dynein complexes expressed using this *DYNC1H1* sequence behaved indistinguishably in *in vitro* motility assays to dynein complexes produced with SNAP-tagged *DYNC1H1* that contained only the H306R mutation. Thus, the E97D mutation appears not to influence dynein motility.

### Baculovirus production and recombinant dynein purification

Baculovirus production and recombinant expression, labelling and purification of individual dynein complexes were performed as described previously (34). Briefly, plasmids encoding SNAP- or Halo-tagged *DYNC1H1* plasmids were recombined with the *pDyn2* plasmid, which encodes human dynein accessory chains that have been codon-optimised for Sf9 cell expression. The accessory chain sequences in *pDyn2* encode DYNC1I2 (DIC2; accession number AF134477), DYNC1LI2 (DLIC2; accession number NM_006141.2), DYNLT1 (Tctex1; accession number NM_006519.2), DYNLL1 (LC8; accession number NM_003746.2) and DYNLRB1 (Robl1; accession number NM_014183.3). Following recombination of the *pDyn1* and *pDyn2* plasmids, the presence of sequences encoding each of the accessory chains was confirmed by PCR. These plasmids were then transposed into the baculovirus genome, followed by another round of PCR-based validation and infection of Sf9 cells.

Dynein complexes were captured from Sf9 cell lysates using IgG Sepharose 6 FastFlow beads (GE Healthcare) in a 2.5 x 10 cm Econo-Column (Bio-Rad) and washed with TEV buffer (50 mM Tris HCl pH 7.4, 148 mM KOAc, 2 mM MgOAc, 1 mM EGTA, 10% (v/v) glycerol, 0.1 mM ATP, 1 mM DTT). SNAP and Halo tagged complexes were labelled at this point with SNAP-Cell TMR-Star (NEB) or Halo-TMR ligand (Promega), respectively (34). Following washing, dynein complexes were released from the beads by TEV protease-mediated cleavage of the LTLT moiety. Dynein complexes were purified by size-exclusion chromatography using a TSKgel G4000SW_XL_ column with a TSKgel SW_XL_ guard column (TOSOH Bioscience) equilibrated in GF150 buffer (25 mM HEPES pH 7.4, 150 mM KCl, 1 mM MgCl_2_, 5 mM DTT, 0.1 mM ATP) using an Ettan LC or AKTA purifier system (GE Healthcare). Each mutant dynein complex eluted with a very similar profile to wild-type dynein. Peak fractions were pooled and concentrated to 0.5–3 mg/ml using Amicon concentrators. To reduce the salt concentration in the dynein preparation, SNAP-dynein complexes were diluted 1:5 in GF50 buffer (GF150 with 50mM KCl) and reconcentrated to the initial protein concentration. This procedure was repeated three times to ensure the final KCl concentration in the buffer was 50 mM. Halo-dynein complexes were dialysed from GF150 to GF50 using Slide-a-lyzer minitubes with a 20 kDa cut-off (ThermoFisher Scientific). All purification and labelling steps were performed at 4°C. Spectrophotometric analysis revealed that all dynein complexes displayed a labelling efficiency of at least 85% of DYNC1H1 monomers (equating to a labelling efficiency of > 97.7% per dimeric dynein complex). Protein aliquots were flash frozen in liquid nitrogen in the presence of 10% (v/v) glycerol and stored at −80°C.

### Purification of dynactin and BICD2N

Native pig dynactin was isolated from fresh pig brains without a microtubule affinity purification step as previously described (55). GFP::BICD2N and SNAP::BICD2N were expressed from baculoviruses in Sf9 cells (34). These proteins contain residues 1–400 of mouse BICD2, which are 95% identical to the equivalent region of the human protein. Purification of GFP::BICD2N and SNAP::BICD2N was performed as described for recombinant dynein except a fluorescent labelling step was not performed before TEV cleavage. We found that labelling of SNAP::BICD2N at this stage could result in inefficient labelling, possibly because the efficient expression of BICD2N meant that the SNAP dye was limiting. Labelling of SNAP::BICD2N was therefore performed after the protein was purified and its concentration measured. The labelling reaction included 4.5 μM of SNAP::BICD2N and 45 μM SNAP-Surface Alexa647 (NEB) and was performed at 4°C for 4h. Excess dye was removed by washing on a PD-10 Sephadex column (GE Healthcare), followed by protein purification by size exclusion chromatography. This procedure resulted in 80% of SNAP::BICD2N monomers being labelled, equating to a labelling efficiency of 96% of dimers. Purified dynactin and BICD2N complexes were stored in GF50 at -80°C in the presence of 10% glycerol.

### SDS-PAGE

SDS-PAGE was performed using Novex 4–12% Bis-Tris precast gels in MES buffer (Life Technologies). Gels were stained with Coomassie Instant Blue (Expedeon) and imaged using a Gel Doc XR+ system with Image Lab 4.0 software (Bio-Rad). Protein concentrations were measured using Quick Start Bradford dye (Bio-Rad) on an Ultrospec 2100 Pro spectrophotometer (Amersham).

### TIRF microscopy

All microscopy was performed at 24–25°C with a Nikon TIRF system equipped with a 100x Apo TIRF objective (1.49 N.A. oil immersion) and the following lasers: 150 mW 488 nm, 150 mW 561 nm (both Coherent Sapphire) and 100 mW 641 nm (Coherent Cube). A back illuminated EMCCD camera (iXon^EM^+ DU-897E, Andor, UK) controlled with μManager software (http://micro-manager.org/wiki/Micro-Manager) was used for image acquisition. The size of each pixel was 105 nm × 105 nm.

### Microtubule gliding assays

Microtubule gliding assays were performed with GMPCPP-stabilised microtubules as described (34). Briefly, flow chambers were passivated with 5% (w/v) pluronic F-127 dissolved in water for 5 min and washed with GF150 buffer. 300 nM of SNAP-dynein or Halo-dynein was diluted in GF50 and injected into the chamber, followed by a 10 min incubation on ice. Unbound motors were removed by washing with GF50, followed by two washes with gliding assay buffer (30 mM HEPES/KOH, 5 mM MgSO_4_, 1 mM DTT, 1 mM EGTA, 40 μM taxol, 1 mg/ml α-casein (Sigma), 2.5mM ATP, pH 7.0). The flow chamber was allowed to warm up to room temperature and a solution injected that contained HiLyte 488-labelled microtubules, 2.5 mM ATP and oxygen scavenging system (1.25 μM glucose oxidase, 140 nM catalase, 71 mM 2-mercaptoethanol, and 24.9 mM glucose) in gliding assay buffer. Microtubules were immediately visualised by TIRF microscopy, with images acquired at 1 s time intervals with 100 ms exposure times. Microtubule velocities were determined by manual analysis of kymographs in Fiji (http://fiji.sc/Fiji). The mean gliding velocity per chamber was determined by subjecting histograms of microtubule velocities to a Gaussian fit.

### Dynein-dynactin-BICD2N motility assays

Assays for processive movement of dynein-dynactin-BICD2N complexes along microtubules were also performed as described previously (34). Briefly, flow chambers were constructed using glass surfaces functionalised with biotinylated poly(ethylene-glycol) (PEG) (67) and counter glass surfaces functionalised with poly(L-lysine)-[g]-PEG (68). GMPCPP-stabilised microtubules labelled with HiLyte488 and biotin were adhered to the glass surface via streptavidin that had been prebound to the biotin-PEG (69). TMR-dynein was incubated with dynactin and BICD2N in motility buffer (30 mM HEPES/KOH, 5 mM MgSO_4_, 1 mM DTT, 1 mM EGTA, 40 μM taxol, 1 mg/ml α-casein (Sigma), 2.5 mM ATP, 50 mM KCl, pH 7.0) for 5 min on ice. The molar ratio in the assembly mix was 1 dynein dimer: 2 dynactin complexes: 20 BICD2N dimers. In the experiments where no dynactin was present (Supplementary Figure 6), dynein and SNAP::BICD2N complexes were mixed at a molar ratio of 1:100. Final dynein concentrations in assembly mixes were typically ˜25 nM. Assembly mixes were typically diluted one to five in motility buffer that was supplemented with ATP (to give a final concentration of 2.5 mM) and oxygen scavenging system and injected into the flow chamber. Complexes were visualised by TIRF microscopy with sequential capture of TMR and Alexa647 signals (200 ms exposure plus 119 ms acquisition per channel), except for the experiments documented in Supplementary Figures 1, 3 and 9, in which only the TMR channel was captured (200 ms exposure plus 36 ms image acquisition). Each chamber in an experimental series used an independent assembly mix.

Kymographs were generated from image series using Fiji software. Typically, 15 microtubules were analysed per chamber. Colocalisation between TMR-dynein and Alexa647-BICD2N was assessed manually in dual colour kymographs. In these and other analyses, only those complexes that associated with a microtubule for ≥ 2.0 s were included. We observed very few instances in which TMR or Alexa647 signals disappeared during processive movements of dual-coloured dynein-dynactin-BICD2N complexes, indicating that photobleaching events during the period of image acquisition were rare. The population of processive, static and diffusive complexes were counted manually as described previously (34). Run lengths and velocities of dynein-dynactin-BICD2N complexes were determined from manual analysis of kymographs. A run was defined as a bout of motion of a unidirectional TMR-dynein that could be terminated by detachment from the microtubule or a pause ≥ 2 s. As noted previously (34), ˜20% of dynein-dynactin-BICD2N complexes switched velocities during a single run. Mean velocity was therefore calculated from individual velocity segments. For run length analysis, a 20-μm segment of each microtubule was selected in Fiji. Chambers in which there were fewer than 10 microtubules ≥ 20-μm-long were not used for run length analysis. Mean run lengths or velocities per chamber were determined by subjecting histograms to exponential or Gaussian fits, respectively.

### Statistics

Prism 6 (Graphpad) was used for data plotting, curving fitting and statistical evaluations. Statistical tests are described in the figure legends.

### Structural analysis

Homology models of human DYNC1H1 were generated using Phyre2 (70) and displayed using PyMOL (https://www.pymol.org/).

## ACKNOWLEDGEMENTS

We thank all members of the Bullock and Carter labs for advice and encouragement. We are particularly grateful to Janina Baumbach, Carly Dix, Helen Foster and Linas Urnavicius for assistance with protein purification or advice. We also thank Alexander Rossor (UCL Institute of Neurology), Mark McClintock and David Garcia (both MRC-LMB) for valuable discussions and comments on the manuscript, and Fillip Port (DKFZ, Heidelberg) for the preprint template. This work was supported by the UK Medical Research Council (file reference numbers MC_U105178790 and MC_UP_A025_1011 for S.L.B. and A.P.C., respectively), the Boehringer Ingelheim Fonds (PhD Fellowship to H. T. H.), the Lister Institute of Preventive Medicine (Research Prize to S.L.B.), the Wellcome Trust (New Investigator Award WT100387 to A.P.C.), EMBO (Young Investigator Award to A.P.C.) and the European Commission (Marie Curie Intra European Fellowship to M.A.S.).

## AUTHOR CONTRIBUTIONS

All authors contributed to the study design and interpretation of the data; H. T. H. performed the majority of experimental work, and evaluated the data; M.A.S. performed several experiments with H.T.H; A.P.C. introduced mutations into *DYNC1H1* constructs. H.T.H., A.P.C. and S.L.B. wrote the manuscript. A.P.C. and S.L.B. supervised the project.

